# KCNQ2/3 regulates efferent mediated slow excitation of vestibular afferents in mammals

**DOI:** 10.1101/2023.12.30.573731

**Authors:** Anjali K. Sinha, Choongheon Lee, Joseph C. Holt

## Abstract

Primary vestibular afferents transmit information from hair cells about head position and movement to the CNS, which is critical for maintaining balance, gaze stability and spatial navigation. The CNS, in turn, modulates hair cells and afferents via the efferent vestibular system (EVS) and its activation of several cholinergic signaling mechanisms. Electrical stimulation of EVS neurons gives rise to three kinetically– and mechanistically-distinct afferent responses including a slow excitation, a fast excitation, and a fast inhibition. EVS-mediated slow excitation is attributed to odd-numbered muscarinic acetylcholine receptors (mAChRs) on the afferent whose activation leads to the closure of a potassium conductance and increased afferent discharge.

Likely effector candidates include low-threshold, voltage-gated potassium channels belonging to the KCNQ (Kv7.X) family, which are involved in neuronal excitability across the nervous system and are subject to mAChR modulation. Specifically, KCNQ2/3 heteromeric channels may be the molecular correlates for the M-current, a potassium current that is blocked following the activation of odd-numbered mAChRs. To this end, multiple members of the KCNQ channel family, including KCNQ2 and KCNQ3, are localized to several microdomains within vestibular afferent endings, where they influence afferent excitability and could be targeted by EVS neurons. Additionally, the relative expression of KCNQ subunits appears to vary across the sensory epithelia and among different afferent types. However, it is unclear which KCNQ channel subunits are targeted by mAChR activation and whether that also varies among different afferent classes. Here we show that EVS-mediated slow excitation is blocked and enhanced by the non-selective KCNQ channel blocker XE991 and opener retigabine, respectively. Using KCNQ subunit-selective drugs, we observed that a KCNQ2 blocker blocks the slow response in irregular afferents, while a KCNQ2/3 opener enhances slow responses in regular afferents. The KCNQ2 blockers did not appear to affect resting afferent discharge rates, while KCNQ2/3 or KCNQ2/4 openers decreased afferent excitability. Here, we show pharmacological evidence that KCNQ2/3 subunits are likely targeted by mAChR activation in mammalian vestibular afferents. Additionally, we show that KCNQ3 KO mice have altered resting discharge rate as well as EVS-mediated slow response. These data together suggest that KCNQ channels play a role in slow response and discharge rate of vestibular afferents, which can be modulated by EVS in mammals.

## Introduction

The vestibular system plays a pivotal role in our sense of balance, orientation, and spatial navigation. Mechanosensitive hair cells (HCs), present among vestibular end-organs in the inner ear, detect both the position and movement of the head. HCs transmit this information to the CNS by way of primary vestibular afferent neurons. The CNS, in turn, can modulate the periphery using the efferent vestibular system (EVS). Vestibular hair cells can be classified into two types: round flask-shaped type I hair cells (HCs) and cylindrical-shaped type II hair cells (HCs). These two types of HCs are synapsed onto by 3 types of primary vestibular afferents: 1. regular firing bouton afferents that synapse onto type II HCs, 2. irregular firing calyx-only afferent that form chalice shaped synapses onto type I HCs; and 3. dimorphic afferents, that synapse onto both type I and type II HC, and vary in discharge regularity from regular (CV*<0.1) to irregular (CV*>0.1) ((Baird et al., 1988); (Fernandez et al., 1988)). These three types of afferents occupy distinct region across the vestibular sensory neuroepithelia, from regular firing afferents in the periphery to irregular firing afferents towards the center of the sensory neuroepithelia.

The CNS, in turn, can modulate the periphery using efferent vestibular system (EVS). EVS neurons originate bilaterally in the brainstem e-group nuclei and extend their axons out with cranial nerve VIII to innervate vestibular end organs in both ears (Gacek & Lyon, 1974; Goldberg & Fernandez, 1980; Jordan et al., 2013; Leijon & Magnusson, 2014; Lorincz et al., 2021; Mathews et al., 2015; Perachio & Kevetter, 1989; Purcell & Perachio, 1997; Warr, 1975). EVS neurons are primarily cholinergic (Bridgeman et al., 1996; Jordan et al., 2015; Kong et al., 1998; Meredith & Roberts, 1987; Pujol et al., 2014) and form extensive varicosities that synapse directly onto type II HCs and afferents throughout the sensory neuroepithelium. EVS innervation of type I hair cells is rare given they are completely enveloped by the calyx afferent ending.

Electrical stimulation of the EVS neurons, in mammals, gives rise to at least three kinetically– and mechanistically-distinct responses including a fast excitation, slow excitation and fast inhibition, so named given their time to peak and duration ((Goldberg & Fernandez, 1980), (Rossi et al., 1980), (Highstein & Baker, 1985), (Brichta & Goldberg, 2000); (Marlinski et al., 2004), (Chagnaud et al., 2015), (Schneider et al., 2021)). Not surprisingly, fast excitation and inhibition are mediated by activation of two distinct nicotinic receptors on afferents and hair cells, respectively, whereas slow excitation, which is the focus of this study, is attributed to the activation of muscarinic acetylcholine receptors (mAChR) on afferent neurons ((Holt et al., 2015), (Holt et al., 2017), (Schneider et al., 2021), (Lee et al., 2021)).

Activation of mAChRs can lead to a number of diverse actions as a function of downstream signaling pathways and effector targets. Of interest to our EVS work, activation of odd-numbered mAChRs (M1/M3/M5) in neurons can close a K+ current called the M-current that gives rise to depolarization and enhanced excitability ((Brown & Adams, 1980) (Marrion, 1997), (Bernheim et al., 1992), also see review (Brown & Yu, 2000)). It was later discovered that the M-current is mediated by members of the KCNQ (Kv7) potassium channel family which are comprised of five distinct subunit subtypes (Kv7.1-Kv7.5) ((Wang et al., 1998); (Suh & Hille, 2002)). A total of four subunits are needed to form a functional K+ channel and different combinations of Kv7.2-Kv7.5 subunits are thought to form K+ channels that can be modulated by mAChR activation (Suh & Hille, 2002). Pore-forming KCNQ channels can be homomeric (containing only one of the KCNQ2-5 subunits) or heteromeric with each channel consisting of multiple KCNQ subunits (See review (Jentsch, 2000); (Delmas & Brown, 2005)). KCNQ2/3, KCNQ2/5, KCNQ3/4, KCNQ3/5 and KCNQ4/5 have been found to form heteromeric channels ((Wang et al., 1998), (Lerche et al., 2000), (Schroeder et al., 2000), (Søgaard et al., 2006), (Bal et al., 2008); (Soh et al., 2022)). Ether-a-go-go-related gene (ERG) (Kv11.1) channel family, particularly erg1/2 heteromeric channels, have also been proposed to underlie some M-currents ((Meves et al., 1999), (Selyanko et al., 2000)). Both ERG and KCNQ (1-5) channels are present in the mammalian vestibular periphery ((Kharkovets et al., 2000), (Hurley et al., 2006), (Lysakowski et al., 2011), (Spitzmaul et al., 2013)). ERG channels are present on both HCs and afferents ((Hurley et al., 2006), (Lysakowski et al., 2011)). KCNQ4 is present on type II HCs and the immature type I HC ((Hurley et al., 2006), (Lysakowski et al., 2011)). In the adult, expression of KCNQ4 decreases in mature type I HCs but it is high out on calyx afferent endings (Hurley et al., 2006). KCNQ4 has higher expression on the inner surface of the calyx ending, facing the type I HCs compared to the outer surface (Lysakowski et al., 2011). Expression of KCNQ3 is higher in the peripheral crista compared to the central zone. KCNQ2 and KCNQ5 are also expressed on calyx bearing afferents but they are expressed at lower levels compared to KCNQ3 and KCNQ4 (Lysakowski et al., 2011). KCNQ1 subunit is not present on the vestibular afferents but it is present on the apical membrane of the marginal cells of the stria vascularis and dark cells in the vestibule where it maintains the endolymphatic potassium concentration (Wangemann, 1995).

In vestibular afferents, the outward K+ current, I_K,M_ is thought to be an M-current ((Hurley et al., 2006), (Rennie et al., 2001), (Perez et al., 2009)). This M-current can be blocked by KCNQ antagonists but not ERG channel blockers ((Perez et al., 2009), (Perez et al., 2010), (Holt et al., 2017), (Ramakrishna et al., 2021), (Bronson & Kalluri, 2023)). Holt et al., (Holt et al., 2017) showed, in turtle, that EVS-mediated slow excitation is mediated by mAChRs and KCNQ channels as it can be blocked by both XE991, a KCNQ channel blocker, as well as the general mAChR blocker, atropine. Patch-clamp studies from mammalian vestibular neurons also showed that mAChRs activation closes the same current as the KCNQ channels ((Ramakrishna et al., 2021), (Bronson & Kalluri, 2023))). While our previous work in mice showed that M3mAChR selective antagonists can block EVS-mediated slow excitation, it has not been shown in an *in vivo* preparation if KCNQ channels are involved in EVS-mediated slow excitation. Additionally, which KCNQ subunits might be utilized for generation of the EVS-mediated slow excitation has not been explored.

In this study, we evaluate the contribution of various KCNQ channel subunits on EVS-mediated slow excitation and the general excitability of primary vestibular afferents in mice. We found that the different KCNQ subunits differentially regulate EVS-mediated slow excitation in regular vs irregular vestibular afferents. While KCNQ2/3 subunits appear to be the main target of EVS-mediated slow excitation in most units, KCNQ3/4 subunits also appear to play critical roles in the maintenance of resting afferent discharge rate.

## Materials and Methods

### Animals

All animal procedures were performed in accordance with NIH’s Guide for Care and Use of Laboratory Animals and approved by the University Committee for Animal Resources (UCAR) at the University of Rochester Medical Center (URMC). C57BL/6J WT mice of both sexes were obtained from Jackson Laboratory and housed in a standard 12-h light:dark cycle with free access to food and water. Mice aged 49-180days, weighing 20-35g were used for the experiments. *Transgenic mice*: KCNQ3 knockout (KO) mice were obtained from Dr. Thomas J. Jentsch at Leibniz-Forschungsinstitut für Molekulare Pharmakologie (FMP) and Max-Delbrück-Centrum für Molekulare Medizin (MDC), Berlin, Germany and bred in house within the URMC vivarium. Here they were also housed in a standard 12-h light:dark cycle with free access to food and water Experiments were performed on mice weighing 20-35g and ages from 49-180 days.

### Animal preparation

C57BL/6J mice were housed in 1-way rooms in the URMC vivarium. Mice were anesthetized with IP urethane (1.6g/kg) and xylazine (20mg/kg). In Figure 5, mice were anesthetized with tribromoethanol (TBE aka Avertin) at 250mg/kg. A tracheotomy was performed for the intubation of PTFE tubing (1.5 cm long, 1mm diam.). The intubation was secured with surgical thread ligatures (Moldestad et al., 2009) and connected to a mechanical ventilator set at 100 bpm (model 683, Harvard Apparatus). A closed-loop homeothermic monitoring system was used to maintain the body temperature. Heart rate was monitored using a 3-lead EKG and a DC amplifier (Warner Instrument, Holliston, MA). Stereotaxic apparatus was used to secure the head of the mice before starting the surgery. After removal of the dorsal scalp and posterior neck muscles, sections of the rear skull were removed with micro-rongeurs to expose the cerebellum and inferior colliculi. The right cerebellar hemisphere, flocculus, and parafloculus were aspirated to expose the arcuate eminence and the common crus. The cerebellar vermis was aspirated using a 5Fr Frazier suction exposing the floor of the 4th ventricle.

#### 3.3.3 **Afferent recordings**

Sharp borosilicate microelectrode (BF150-86-10, Sutter Instruments) with impedance 60-120 MOhms was used to record extracellular spike activity from spontaneously discharging vestibular afferents. The microelectrode, filled with 3M KCl, was mounted on a 3-axis micromanipulator and slowly advanced into the superior division of CNVIII just as it exits the otic capsule. After connecting to a preamplifier headstage (Bio-medical Engineering, Thornwood, NY), microelectrode capacitance was neutralized by driving the shield of the input cable, and electrode resistance was balanced using dial settings on the amplifier.

### Efferent stimulation

To stimulate EVS neurons in the brainstem, we used a linear 4-lead platinum-iridium array. The stimulating electrode array was placed into 4th ventricle at the midline with the first electrode lead located just caudal to the facial colliculi (Schneider et al., 2021). Efferent stimulation was delivered from a stimulus isolator (model A360; World Precision Instruments, Sarasota, FL, United States). In-home spike2 script was used to manage the timing of the stimulation via a micro1401 interface (Cambridge Electronic Design). Standard EVS stimuli consists of 5-s trains of 100-150µsec constant current shocks was delivered at 333 shocks/s (Brichta & Goldberg, 2000). The shock pulses amplitude was adjusted to determine the threshold (T, 20-50uA) and maximum current (40-250 uA) in order to avoid antidromic activation.

### Data acquisition

Data acquisition was managed using in-house Spike 2 scripts with micro1401 interface (Cambridge Electronic Design). Afferent voltage signals were low-pass filtered (1KHz, four-pole Bessel; Wavetek) and sampled at 10 KHz. Spike2 data files were analyzed using IgorPro 6.36 (WaveMetrics). Stimulation artifacts were removed offine by subtracting an averaged shock artifact from each shock stimulus in the raw data.

### Drug Administration

All drugs were administrated in one of two ways: 1) Systemically using standard intraperitoneal (IP) injection or 2) Delivery into middle ear space via Intrabulla administration (IB). IB administration of drugs was done as explained before (Lee et al., 2021, Suzuki et al., 2017). Briefly, for IB injections, an incision was made behind the right pinna and bulla was exposed by removing the muscle above it. A hole was made in the otic bulla using 30-G needle to insert a 100-200um diameter tube. This tube was used to administer drugs to access the middle ear space. The tube was held in place using cyanoacrylate glue.

### Drug used

XE991, E-4031, ICA110381, UCL2077, ML213 and Oxotremorine-M were obtained from Tocris (Tocris Biosciences, Bio-Techne, UK). Retigabine was obtained from Toronto Research Chemicals (Toronto, ON, Canada). Stock solutions for XE-991, E-4031 and Oxotremorine-M were prepared in water and diluted to our daily working dilution in saline for systemic administration. Stock solutions and working dilutions for ICA 110381, UCL2077 and ML213 were prepared in 100% DMSO. Unless specified, all drugs were administered systemically through intraperitoneal (IP) route. Drugs in 100% DMSO were systemically administered at 25ul DMSO/10g of mice with maximum of 100ul of DMSO total. Retigabine acts on KCNQ channels by stabilizing the S5-6 domain in the open state accompanied by a hyperpolarizing shift in the voltage activation curve and an increase in deactivation time ((Brown & Passmore, 2009; Wuttke et al., 2005)). ICA 110381 bind to a different site on the KCNQ channel to decrease the input resistance of the cell and shift the resting membrane potential in more hyperpolarized direction (Boehlen et al., 2013). ML213 also shifts the voltage activation curve more negative ((Yu et al., 2011), (Brueggemann et al., 2014)). UCL2077, on the other hand, is cell membrane permeable and inhibits the channel by blocking the pore from the inside ((Barro-Soria et al., 2014)). In Figure 6, UCL2077 was administered through IB route in 100% DMSO. 100% DMSO non-significantly enhances baseline (Sinha et al., 2023). Systemic administration of DMSO did not affect vestibular afferent firing when injected at volume less that 100ul (Lee et al., 2017).

### Vestibular sensory Evoked Potential (VsEP)

Linear VsEP (denoted here as VsEP) is compound action potential from macular vestibular afferents, originating mostly from saccule and utricle, as population response to transient linear acceleration and recorded on the surface of the scalp. For the purpose of this study the first three peaks, P1, N1 and P2 were scored and analyzed. The animal preparation and the VsEP recording to linear acceleration pulses were done according to previously published work by Jones and colleagues (Jones et al., 1999; Jones et al., 2001; Jones et al., 2002). Briefly, animals were anesthetized using IP administration of urethane (1.6g/kg)/xylazine (20mg/kg). A tracheotomy was performed for the intubation of PTFE tubing (1.5 cm long, 1mm diam.), secured with surgical thread ligatures (Moldestad et al., 2009) and connected to a mechanical ventilator set at 100 bpm (model 683, Harvard Apparatus). Animal was placed in supine position on a heated pad to maintain body temperature at37.5 Celsius. using a closed-loop homeothermic monitoring system. Heart rate was monitored using a 3-lead EKG and DC amplifier (Warner Instrument, Holliston, MA). Recording electrodes were placed epidurally on the dorsal scalp (non-inverting electrodes) and behind the right pinna (inverting electrodes) and on the hip (ground). The head was secured to a voltage-controlled mechanical shaker via a head mount. Linear acceleration pulses were generated by using custom software. A linear voltage ramp of 2-ms duration was applied to an electromagnetic shaker (model ET-132-203; LabWorks, Costa Mesa, CA). Stimuli were presented at the rate of 17 pulses/s. Stimulus amplitude was calibrated and monitored using a calibrated accelerometer mounted on the shaker platform. Amplitude was quantified in decibels relative to the reference unit of jerk (specifically dB re: 1.0 g/ms, where 1.0 g=9.8 m/s^2^). Stimulus amplitude ranged from –15 to +6 dB re: 1.0 g/ms. Vestibular stimuli were presented with a binaural forward masker of broadband noise (50-50,000 Hz bandwidth, 90dB SPL). The masker was presented with a free-field speaker. This was done to eliminate the auditory contribution to VsEP data. The electrophysiological signal was amplified (200,000 times; Grass P511) and filtered (0.3-3 kHz). The signal was digitized at the onset of each stimulus (1,024 points at 10us/point). Signal averaging was employed to enhance the signal-to-noise ratio. To do this, separate averages traces of each polarity was collected (128 sweeps each) which was then averaged for 256 sweeps. During the study involving Oxotremorine-M (Oxo-M), after administering Oxo-M intraperitoneally at a concentration of 2 mM, we recorded VsEPs at +6 dB (re: 1.0 g/ms) at 5-minute intervals. Increased salivary secretion occurred due to administration of Oxo-M and was wicked using wipes. Additionally, we determined the VsEP threshold every 10-15 minutes for a duration of 30-40 minutes. Total of 14 controls were used in the analysis of VsEP data. Five (5) controls were collected simultaneously with the KCNQ3 KO mice. 11 controls were used from the previous work (Sinha et al., 2023). There was no significant difference between the two control groups and hence the data from these two experiments were combined for analysis to increase the power of the groups.

### Statistical procedures

The effects of different pharmacological treatments on afferent discharge rate and efferent-mediated slow excitation were assessed using a paired t-test. All data were tested for normality. Paired t-test was also used to compare slow response amplitude between control and KCNQ3 KO mice. Values, expressed as means, SEM, and outcome parameters including p-values, F-statistics, t-statistics, and effect sizes (Cohen’s d) are reported in the text and/or figures. Mixed-model ANOVA with Šídák’s multiple comparison was used to analyze effect of time or stimulus intensity on VsEP amplitude and latency between control and KCNQ3 KO mice. All statistical analyses were done in GraphPad Prism. Values, expressed as means ± SEM, and outcome parameters including p values are reported in the text and figures. For figure 5 and 6 control data from (Sinha et al., 2023) was reused (n=5 animals) for discharge regularity and EVS-mediated slow excitation. Additionally, four (n=4) animals were recorded simultaneously with the KCNQ3 KO mice.

## Results

### M-current through KCNQ channels underlie EVS-mediated slow response

EVS-mediated slow excitation results from the activation of odd-numbered mAChRs on vestibular afferents that subsequently block an M-current ((Holt et al., 2017), (Schneider et al., 2021); (Lee et al., 2021), (Perez et al., 2009), (Ramakrishna et al., 2021)). The M-current may be mediated by either member of the KCNQ channel subfamily (Kv7.2-Kv7.5) ((Wang et al., 1998), (Suh & Hille, 2002)) or the ERG channel family (Erg1-Erg3) ((Meves et al., 1999), (Selyanko et al., 2000)). While Kv7.2-7.5 and ERG2/3 subunits are present on mammalian vestibular afferents (Hurley et al., 2006, Lysakowski et al., 2011), previous vestibular studies have implicated KCNQ channels as the likely target of activation of mAChRs ((Perez et al., 2009), (Holt et al., 2017), (Ramakrishna et al., 2021)). To confirm which potassium channel family is involved in mice, we examined the effects of both ERG and KCNQ blockers on EVS-mediated slow excitation. In Figure 1, we compared EVS-mediated slow excitation of mouse vestibular afferents before and after the administration of either the KCNQ channel blocker, XE991, (Figs. 1A-D) or the ERG channel blocker, E-4031 (Figs. 1E-F). In the control periods, repeated shock trains (333 shocks/s for 5 seconds every 60 seconds) gave rise to a slow excitation characterized by its slow time to peak and extended return to baseline (Figs. 1A, 1E). IP administration of XE991 inhibited the EVS-mediated slow excitation (n=6/7, t(5)=3.098, p=0.0.0269) (slow excitation: n=7/7, 20.27 ± 5.954 vs 6.405 ± 3.418, t(6)=1.940, p=0.1004; student’s paired t-test) (Fig. 1A-B). The baseline discharge rate of afferent also went up in most cases (Fig. 1A) (n=5/7) (baseline: n=7/7, 30.87±12.93 vs 55.76±8.915, t(6)=2.015, p=0.0905, student’s paired t-test) (data not shown). Additionally, local administration via the intrabulla (IB) route, of XE991 (0.5mg/ml) also blocked most of the EVS-mediated slow excitation (20.58 ± 5.04 vs 4.77 ± 1.59, t(6)=2.015, p=0.0137) (Fig. 1C), whereas IB administration of E-4031, at two concentrations, did not have a significant effects on the slow response amplitude (Fig. 1F) (2mg/ml: 10.21 ± 2.35 vs 11.43 ± 3.81, t(5)=0.6074, p=0.5701; 20mg/ml:: 8.21 ± 2.351 vs 5.96 ± 1.49, t(2)=1.222, p=0.3461). Application of XE991 also enhanced the baseline firing rate in most of afferents recorded (p=0.0203) while in one unit, the baseline went down (n=7, 40.16 ± 11.45 vs 49.75 ± 6.13, t(6)=1.050, p=0.331) (Fig. 1A, 1D). This is likely due to depolarization of afferents, as a result of blocking many of the KCNQ channels on the afferents. Local and systemic administration of XE991 had similar effects. This is in line with previous studies where the M-current was blocked by XE991 ((Holt et al., 2017), (Ramakrishna et al., 2021)) but not with E-4031 (Perez et al., 2009).

**Figure 1.**
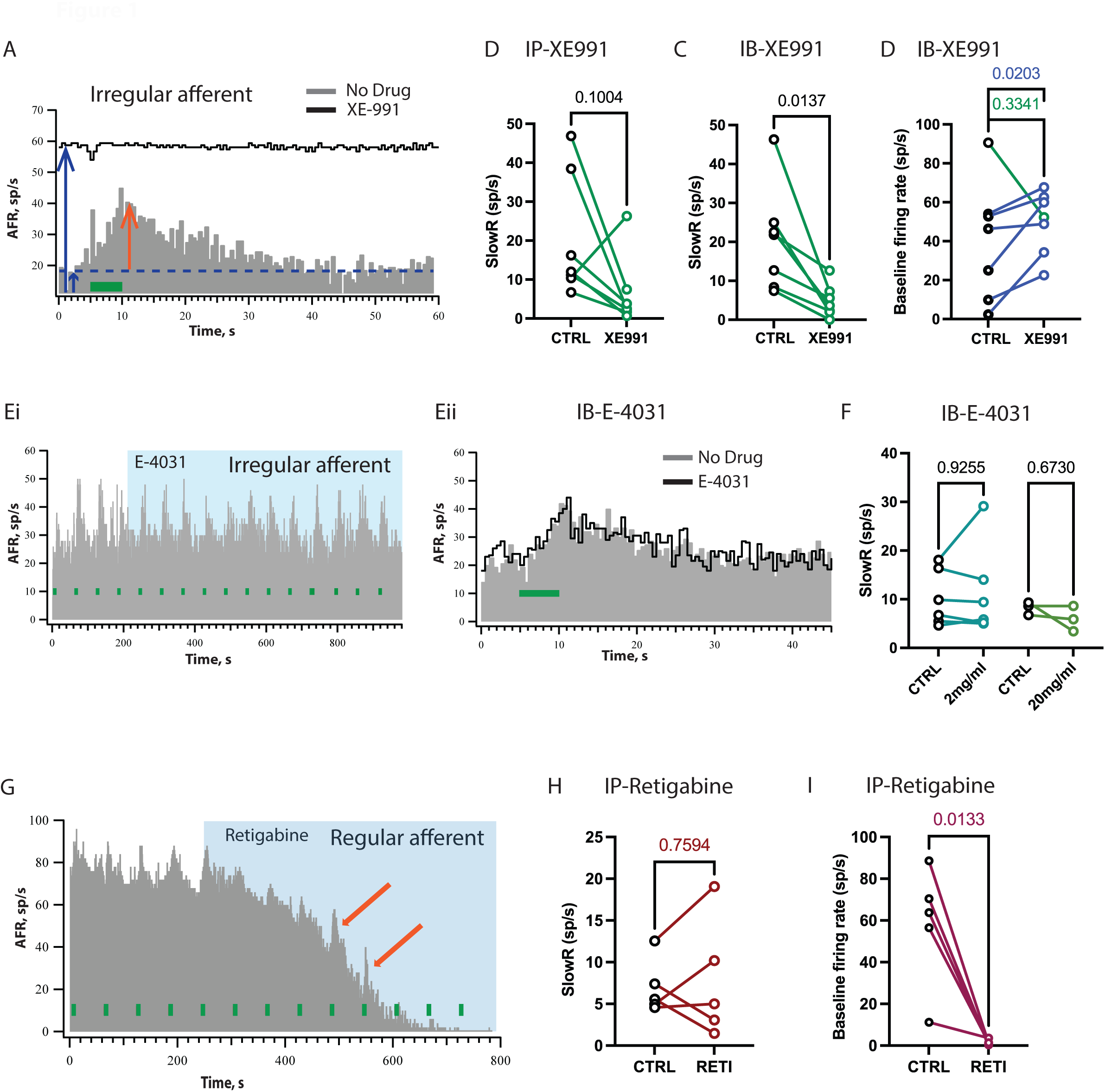
KCNQ underlie EVS-mediated slow excitation. A, Average rate histograms of pre-(grey) and post-(black) reveal KCNQ channel antagonist, XE991(5mg/kg IP) blocks the slow response and enhances baseline (green bars, 333/s for 5s every 60s). Blue dashed line represents afferent baseline discharge rate. Blue arrows indicate the amplitude of baseline firing rate in both pre– and post-drug condition. Orange arrows indicate amplitude of slow response (SlowR). B,C, Summary data for effect of IB administration of XE991 on slow response (SlowR). D, baseline firing rate (student’s paired t-test). Ei, Continuous rate histograms before and after E-4031, ERG channel antagonist, Eii, average rate histograms show IB administration of the ERG channel antagonist, E-4031 (2mg/ml or 20mg/ml) does not block the slow excitation (n=8). F, Average rate histograms of pre-(grey) and post-(black) IB administration of E-4031 (2mg/ml, student’s paired t-test). G, Continuous rate histogram reveals that IP administration of the KCNQ agonist, retigabine, enhances slow response and leads to drop in baseline. Orange arrows show the units with enhancement in slow response post-retigabine H, Summary data for effect of retigabine on slow response. I, Summary data for effect of retigabine on baseline firing rate.

To further explore a role for KCNQ channels, we also examined the effects of the general KCNQ channel opener, retigabine (Tatulian et al., 2001) on EVS-mediated slow excitation in mouse vestibular afferents. By targeting the same KCNQ channels, we expected that retigabine would produce opposite effects of those seen with XE991 (Tatulian et al., 2001). An increase in the number of open KCNQ channels on the afferent should lead to decrease in afferent excitability and resting discharge. Additionally, if some fraction of the open KCNQ channels can also be engaged by EVS stimulation, one might expect an enhancement in the amplitude of EVS-mediated excitation provided those newly opened KCNQ channels can now be closed by mAChR activation. As predicted, IP application of retigabine (20mg/kg) produced a significant decrease in afferent baseline firing rate (Figs. 1G, 1I) (58.11±12.88 vs 1.4±0.57, t(4)=4.239, p=0.0133, student’s paired t-test). Interestingly, during the decrease in the afferent discharge rate within 3-6 mins post-IP administration of retigabine, EVS-mediated slow excitation grew in amplitude compared to the pre-drug condition. We saw this enhancement in 3 of 5 afferent recordings (n=5, 7.016±1.466 vs 7.76±3.186, t(4)=0.3280, p=0.7594, student’s paired t-test) (Fig. 1H). Like the data with XE991, there seems to be two populations of responders. At later time points (7-10 minutes), there was a near complete collapse in the afferent’s firing rate presumably as a result of the continued opening of additional KCNQ channels by retigabine. Collectively, these data when combined with the lack of effect with the ERG blocker, E4031, suggests that members of the KCNQ channel family underlie the M-current whose closure by mAChRs gives rise to EVS-mediated slow excitation of mouse vestibular afferents.

### KCNQ2/3 opener enhances slow response amplitude in regular firing afferents

Although we now know that members of the KCNQ subfamily regulate the M-current in mouse vestibular afferents, it is not immediately clear which KCNQ channel subunits are involved. KCNQ2-5 subunits are present on mammalian calyx-bearing afferents and appear to be differentially expressed within different microdomains of the calyx ending which may also vary depending on that calyx ending’s position in the sensory neuroepithelia (Hurley et al., 2006), (Lysakowski et al., 2011)). To explore which KCNQ channel subunits might contribute to EVS-mediated slow excitation and general afferent excitability, we relied on the use of several subunit-selective KCNQ openers and blockers. Because the distribution of different KCNQ subunits also vary among regular and irregular discharging afferents, we present the pharmacological data in regular and irregular afferents separately.

We first systemically administered ICA110381 (ICA), a KCNQ2/3 opener, at 5mg/kg (Boehlen et al., 2013). KCNQ2 is present on inner surface of calyx endings and KCNQ3 is present on the outer surface of dimorphic afferents and extends up to the heminode (Lysakowski et al., 2011). IP administration of ICA showed similar effects on afferent neuron firing as retigabine. Administration of ICA led to decrease in afferent firing rate in all afferent units recorded (Figs. 2A-F) (Regular: 67.98±7.04 vs 1.68±1.17, t(9)=9.512, p<0.0001; Irregular: 34.87±12.30 vs 0.15±0.14, t(5)=2.806, p=0.037, student’s paired t-test) (Regular: Figs. 2A-C; Irregular: Figs. 2D-F). However, the KCNQ2/3 opener had differential effect on the EVS-mediated slow response in regular (CV*<0.1) vs irregular firing afferents (CV*>0.1). In regular afferents, we saw enhancement of EVS-mediated slow response amplitude (n=8/10 of regular afferents), similar to retigabine. The exact time of enhancement of slow response varied based on when the drug reached the inner ear following the IP administration (Figs. 2A-B). The relative timing of drug reaching the inner ear can be inferred from when we start to see the decrease in afferent firing rate (Figs. 2A, B). To account for timing differences in the arrival of ICA, we aligned the peak enhancement in EVS-mediated slow excitation from each affected afferent (Figs. 2G-H). Here, we can clearly see that the mean EVS-mediated slow response amplitude more than doubles in size during ICA application (Figs. 2A, G) followed by a collapse of EVS-mediated slow response, due to disruption of the afferent resting discharge rate. This indicates that as the concentration of ICA increased at the synapse, more KCNQ channels are opened. And as more KCNQ channels opened, more channels on the afferents were also available for closure by mAChR activation. This led to a gradual increase in the slow response amplitude (pre-drug slow response amplitude vs peak slow response amplitude: n=8/10 of regular units: 9.76±1.19 vs 23.52±4.71, t(7)=3.025, p=0.0192; n=10/10 of regular units: 9.773±0.9424 vs 18.82±4.868, t(9)=1.897, p=0.0903, student’s paired t-test).

**Figure 2.**
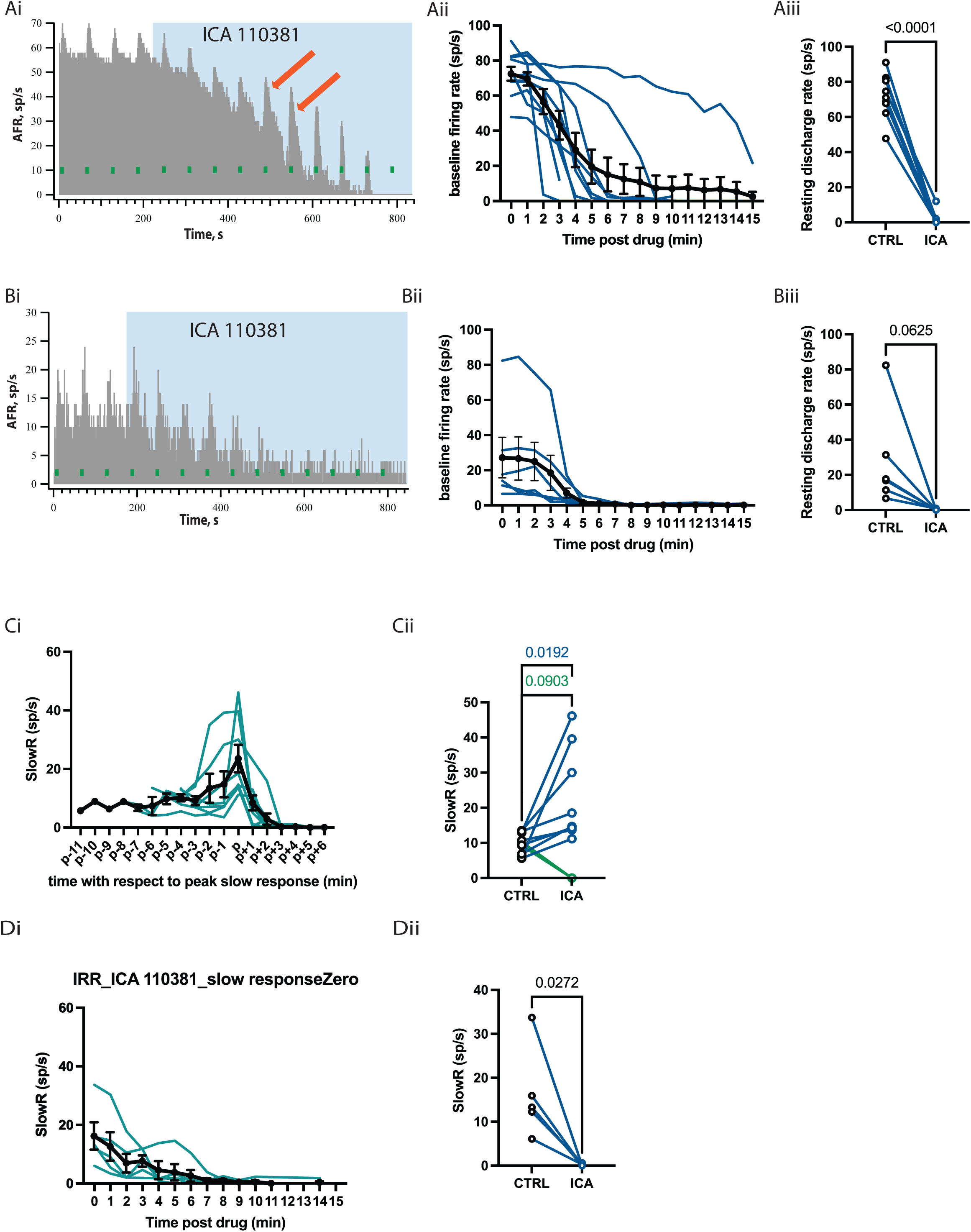
KCNQ2/3 opener enhances EVS-mediated slow response in regular firing afferents. A, Continuous rate histograms of regular afferents (CV*<0.1). D, and irregular firing afferents (CV*>0.1). Continuous rate histogram shows changes in afferent firing rate (AFR) during midline efferent stimulation (green bars, 333/s for 5s every 60s). B, E, Individual traces of continuous response histogram of baseline firing rate of regular (CV*<0.1, B) and irregular (CV*>0.1, E) firing afferents post administration of ICA110381, KCNQ2/3 opener, at 5mg/kg. C, F, Baseline amplitudes are plotted for the regular (C) and irregular (F) afferents in the panel as pre– and post-ICA110381 administration. G, H. Mean slow excitation, averaged from a 1-s block at t = 6–7 s, post administration of ICA110381. G, Peak enhancement (p) in slow excitation is aligned for n=8/10 regular units (blue, individual traces; black, average of 8 regular units). G, Slow response amplitude are plotted for all the regular (H) units pre– and post-ICA110381 administration (dark blue, unit enhancement in slow response and p-value for traces only with enhancement post-drug (8/10); light blue lines, regular units with no enhancement in slow response amplitude, p-value for all the regular unit, n=10/10) (student’s paired t-test). I, Slow response trace post administration of ICA110381 for all irregular units (n=5) (blue trace, individual lines; black trace, average trace). J, Slow response amplitude are plotted for all the irregular units pre– and post-ICA110381 administration (student’s paired t-test).

In contrast, however, we did not see an enhancement in the EVS-mediated slow excitation in any of the irregular firing afferents we recorded. What we saw instead was a gradual decrease in the mean amplitude of EVS-mediated slow excitation that generally followed the decrease in afferent firing rate (Figs. 2I, 2J) (16.25±4.660 vs 0.1844±0.1254, t(4)=3.403, p=0.0272, student’s paired t-test). This can happen for 2 reasons. Firstly, the ICA concentration at the synapse was always much higher compared to number of KCNQ2/3 channels and access to different regions of the neuroepithelium. The stroma is notably thicker in peripheral regions and may also rise to delayed drug entry. As such, this could lead to collapse of the afferent firing rate before we could record any possible enhancement in the amplitude of EVS-mediated excitation. The second possibility is that KCNQ2/3 channels do not contribute in the same way to EVS-mediated slow excitation in irregular afferents beyond general effects on afferent excitability.

### KCNQ2 blocker inhibits slow response but does not affect resting discharge rate

Next, we used UCL2077 (UCL), a KCNQ1/2 blocker (Shah et al., 2001, Soh et al., 2010, Barro-Soria et al., 2014). KCNQ1 is present on darks cells in the vestibule and marginal cells in the stria vascularis which is involved in maintaining the endolymphatic potential. Lee et al., (Lee & Jones, 2018) showed that blocking Na+-K+-2Cl-cotransporter (NKCCl), which is needed to maintain K+ secretion from the dark cell to maintain homeostasis, did not alter vestibular sensory evoked potential (VsEPs) but did alter the auditory brainstem response (ABR). VsEP is a compound action potential generated by irregular firing afferent in the macula, in response to a jerk stimulation (Jones, 1992; Jones et al., 2015; Ono et al., 2020). This suggests that although firing of auditory afferents is disrupted by alterations of the endolymphatic potential, vestibular afferents are functional for over an hour post-IP administration of the NKCCl blocker. These observations and the fact that KCNQ1 is not expressed in HCs or vestibular afferents suggesting that we can attribute the effect of UCL on vestibular afferent to the blockade of KCNQ2 channels. KCNQ2 channel subunits are primarily present on the inner surface of calyx endings.

Systemic administration of UCL at 20mg/kg did not affect the baseline firing rate of vestibular afferents (Fig. 3A-F) (Regular: 60.3±7.16 vs 63.59±8.41, t(3)=1.747, p=0.1790; Irregular: 38.71±7.84 vs 35.33±8.90, t(8)=1.335, p=0.2185) (Figs. 3C, 3F). Similar to effects seen after ICA administration, the KCNQ2 blocker UCL also had differential effect on regular and irregular afferents. UCL reduced the EVS-mediated slow response by 66% in irregular firing afferents (Figs. 3I-J) (14.03±2.4 vs 4.73±1.34, t(8)=3.12, p=0.014, Fig. 3J) whereas in regular afferent the slow response was only blocked by 26% (Figs. 3G-H) (9.36±1.03 vs 6.86±0.36, t(3)=2.675, p=0.0754, Fig. 3.3H). This suggests that the KCNQ2 channel plays a larger role in EVS-mediated slow responses in irregular firing afferent compared to regular firing afferents. However, KCNQ2 is not a major contributor of resting discharge rate of vestibular afferents, at least under our recording conditions. Even though it did close KCNQ channels on the irregular vestibular afferents, we did not see any change in baseline firing rate.

**Figure 3.**
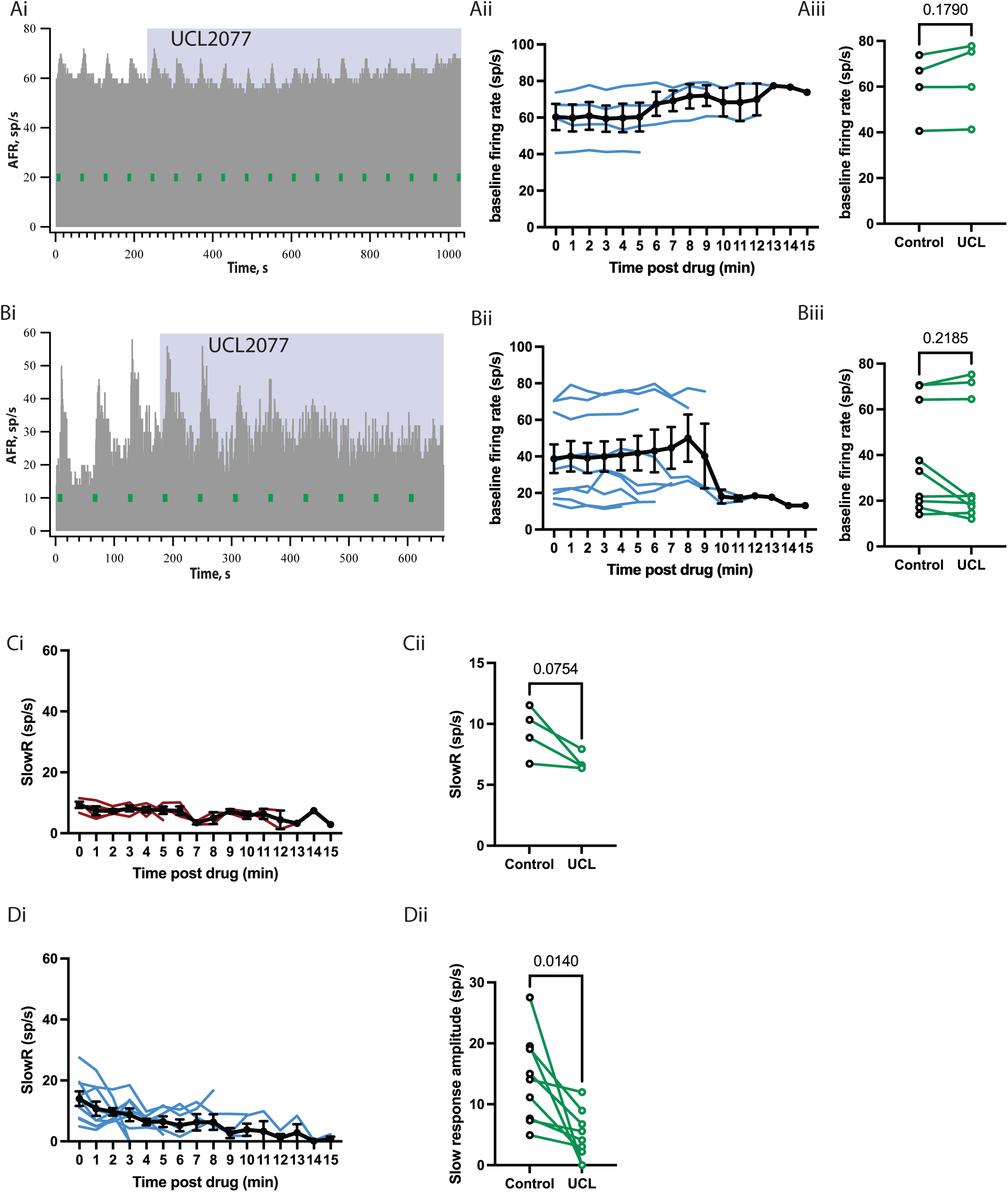
KCNQ2 blocker, UCL2077, blocks EVS-mediated slow response: A, D. Continuous rate histograms of regular afferents (CV*<0.1) (B) and irregular firing afferents (CV*>0.1) (D) shows changes in afferent firing rate (AFR) during midline efferent stimulation (green bars, 333/s for 5s every 60s). B, C, Individual traces of continuous response histogram of baseline firing rate of regular (CV*<0.1, B) and irregular (CV*>0.1, C) firing afferents post administration of UCL2077 (UCL), KCNQ2 blocker, at 20mg/kg. C, F, Baseline amplitude are plotted for the regular (B) and irregular (F) afferents in the panel as pre– and post-UCL administration. G-J, Mean slow excitation, averaged from a 1-s block at t = 6–7s, post administration of UCL. G, Slow response trace post administration of UCL for all irregular units (n=4) (blue trace, individual lines; black trace, average trace). H, Slow response amplitude are plotted for all the regular units pre– and post-UCL administration (student’s paired t-test). I, Slow response trace post administration of UCL for all irregular units (n=9) (blue trace, individual lines; black trace, average trace). J, Slow response amplitude are plotted for all the irregular units pre– and post-UCL administration (student’s paired t-test).

### KCNQ4 contributes to resting discharge rate of the vestibular afferent: heterogenous responses

Of all the available KCNQ subunits expressed in the inner ear, KCNQ4 channels have received the most attention. Mutations in the KCNQ4 gene leads to DFNA2, an autosomal dominant progressive deafness, in humans (Kharkovets et al., 2000) due to slow degeneration of outer HCs (Kharkovets et al., 2006). KCNQ4 KO mice also have a reduced vestibular ocular reflex (VOR) gain (Spitzmaul et al., 2013). Recent studies have also suggested a role for KCNQ4 in non-quantal transmission from HCs to afferents ((Contini et al., 2017), (Contini et al., 2020), (Govindaraju et al., 2023)).

The KCNQ4 subunit is most prominent on the inner surface of calyx endings and to a smaller extent on the outer surface. KCNQ4 is also present in the axon heminode on calyx-only afferents (Lysakowski et al., 2011). Due to the presence of the KCNQ4 subunit at different locations within the calyx ending as well as differential distribution between calyx-only and dimorphic afferents. We anticipated considerable variability in the effects of a KCNQ4 opener on afferent excitability and potentially on EVS-mediated slow excitation. While we can distinguish afferents based on discharge regularity in our preparation, we could not identify whether we were recording from calyx-only afferents and irregular dimorphic afferents.

To explore the role of KCNQ4 channels in EVS-mediated slow excitation and afferent excitability, we used the KCNQ2/4 opener ML213. Systemic administration of ML213 decreased the resting discharge rate in all vestibular afferents recorded (Regular: 67.88±6.38 vs 15.26±8.33, t(10)=5.894, p=0.0002; Irregular: 28.13±5.44 vs 1.48±0.81, t(12)=5.006, p=0.0003, student’s paired t-test). Interestingly, some of the units recorded, had a sudden collapse in resting discharge rate (regular: 3/10; irregular: 1/13) (Fig. 4A) while others demonstrated a gradual decline in resting discharge rate (Figs. 4D-E) (regular: 3/10; irregular: 12/14). Some of the units had a gradual decline in afferent firing rate followed by a sudden collapse of afferent discharge (Fig. 4B) (regular: 4/10; irregular: 0/14). The kinetics of decrease in resting discharge could depend on the location of the KCNQ4 on the calyx ending and the time it takes for the drug to reach that location.

**Figure 4.**
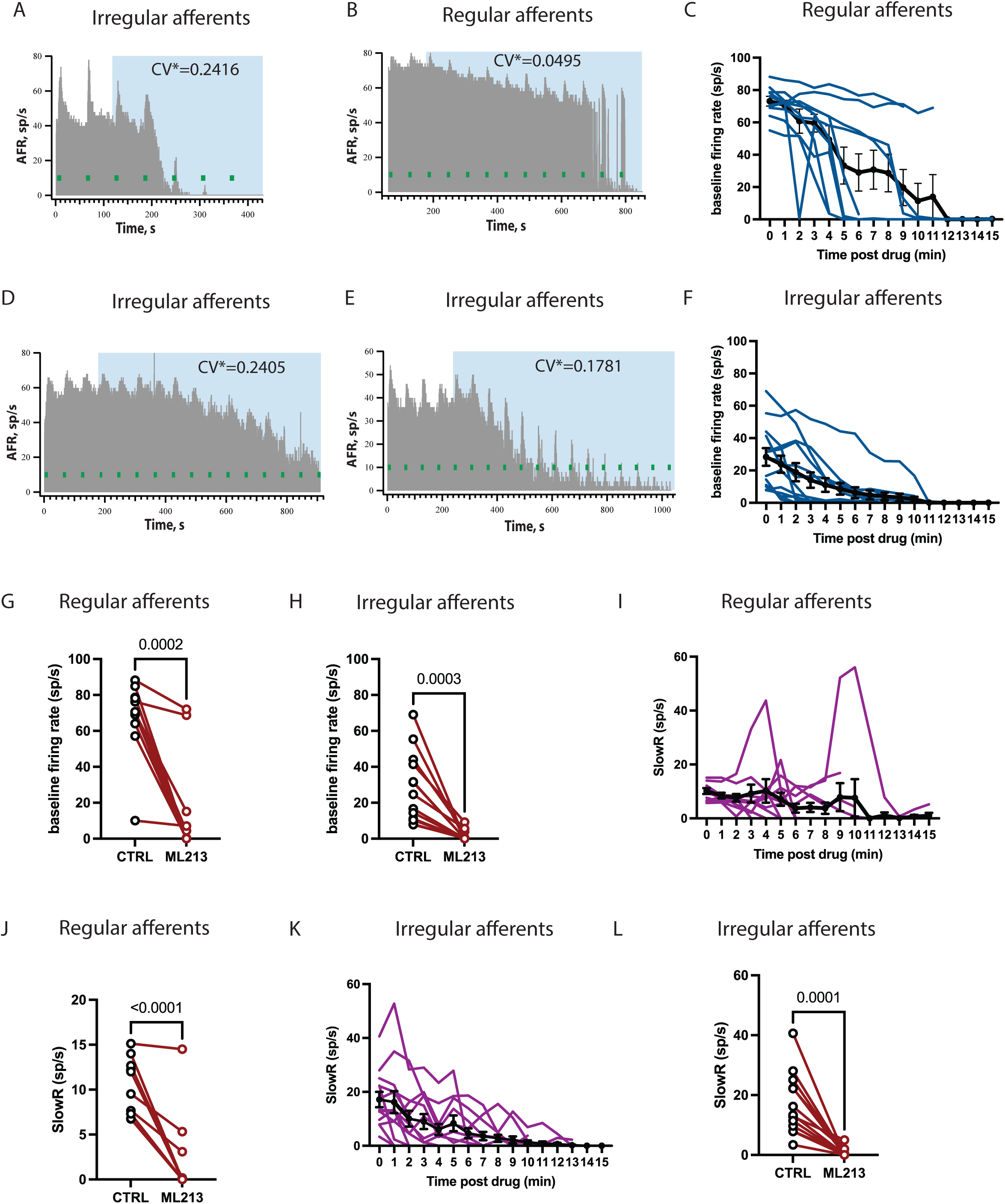
KCNQ2/4 opener, ML213, attenuates baseline firing rate: A, D, Continuous rate histogram of regular (A) and irregular units (D), post-administration of 5mg/kg of ML213. A, Continuous rate histograms of a regular afferent (CV*<0.1), post-administration of 5mg/kg of ML213, with sudden drop in afferent firing rate (AFR). B, Continuous rate histograms of a regular afferent (CV*<0.1), with gradual decrease in baseline followed by sudden drop of AFR. D, Continuous rate histograms of an irregular afferent (CV*>0.1), post-administration of 5mg/kg of ML213, with gradual decrease in baseline with no change in slow response amplitude. E, Continuous rate histograms of an irregular afferent (CV*>0.1), with gradual decrease in AFR with enhancement in slow response (yellow arrow). C, F, Baseline amplitude are plotted for the regular (C) and irregular (F) afferents post-administration of ML213 (red trace, individual units; black trace, average trace). G, H, Baseline amplitude are plotted for the regular (G) and irregular (H) afferents in the panel as pre– and post-administration of ML213 at 5mg/kg. I-L. Mean slow excitation, averaged from a 1-s block at t = 6–7 s, post administration of ML213. I, K, Slow response trace post administration of ML213 for regular (I) and irregular units (K) (purple, individual lines; black trace, average trace). J, L, Slow response amplitude are plotted for all the regular units (J) and irregular units (L), pre– and post-administration of ML213 (student’s paired t-test).

ML213 also had differential effects on EVS-mediated slow excitation. In most of the regular and irregular firing afferents there was a decrease in the EVS-mediated slow response associated with the decrease in resting discharge rate (Figs. 4A, 4D). The slow response was lost with the collapse in afferent firing presumably based on gross changes in excitability (regular: n=8/10, irregular: n=14/14) (Fig. 4J, 4L) (regular: 10.55±0.92 vs 2.10±1.34, t(10)=6.460, p<0.0001; vs irregular: 16.98±2.85 vs 0.73±0.40, t(12)=5.568, p=0.0001). In the case of 5 regular units, there was a delayed increase in the EVS-mediated excitation, after the collapse of afferent discharge rate (Figs. 4B, 4I). In these cases, the enhancement might be an indirect effect of EVS-stimulation. Closure of the KCNQ channels, due to activation of mAChRs, increased the impedance of afferents. This increase in impedance might be enough to allow for the generation of action potential in the afferents by bringing the threshold near the firing potential, leading to resurrection of afferent firing with slow response on top (Figs. 4B, 4I). Additionally, two regular and three irregular units showed enhancement in slow response amplitude with application of ML213 (Figs. 4B, 4E). This enhancement might be attributed to opening of KCNQ2 channel subunits. Alternatively, KCNQ4 might also be involved in slow excitation, but it is difficult to parse out using our preparation. These observations suggest that, at least in our preparation, KCNQ4 is involved in maintaining the resting discharge rate in all vestibular afferent and might also regulate EVS-mediated slow response in some of the afferents.

### KCNQ3 KO mice have altered resting discharge rate distribution of vestibular afferents

To further characterize the role of KCNQ3 in maintaining homeostasis of vestibular afferent excitability and in EVS-mediated slow excitation, we explored both in KCNQ3 knockout (KCNQ3 KO) mice. We first characterized the distribution of afferent resting discharge rates against CV*. The distribution of resting discharge rates was altered in KCNQ3 KO mice (n=54units/8 animals) compared to control (n=76units/9 animals). The KCNQ3 KO mice had fewer high firing (>90 sp/s) afferents compared to control (Figs. 5A, 5B). Moreover, KCNQ3 KO mice had a distribution shift in CV* compared to control (Fig. 5C control vs KCNQ3 KO: n=9 vs 8 animals, see method) particularly as an increase in percentage of units with intermediate discharge rate (40< Resting discharge rate<60) (Fig. 5D, control vs KCNQ3 KO: n=9 vs 8 animals, see method). Altogether, these data that the discharge properties of vestibular afferents in KCNQ3 KO have been altered.

**Figure 5.**
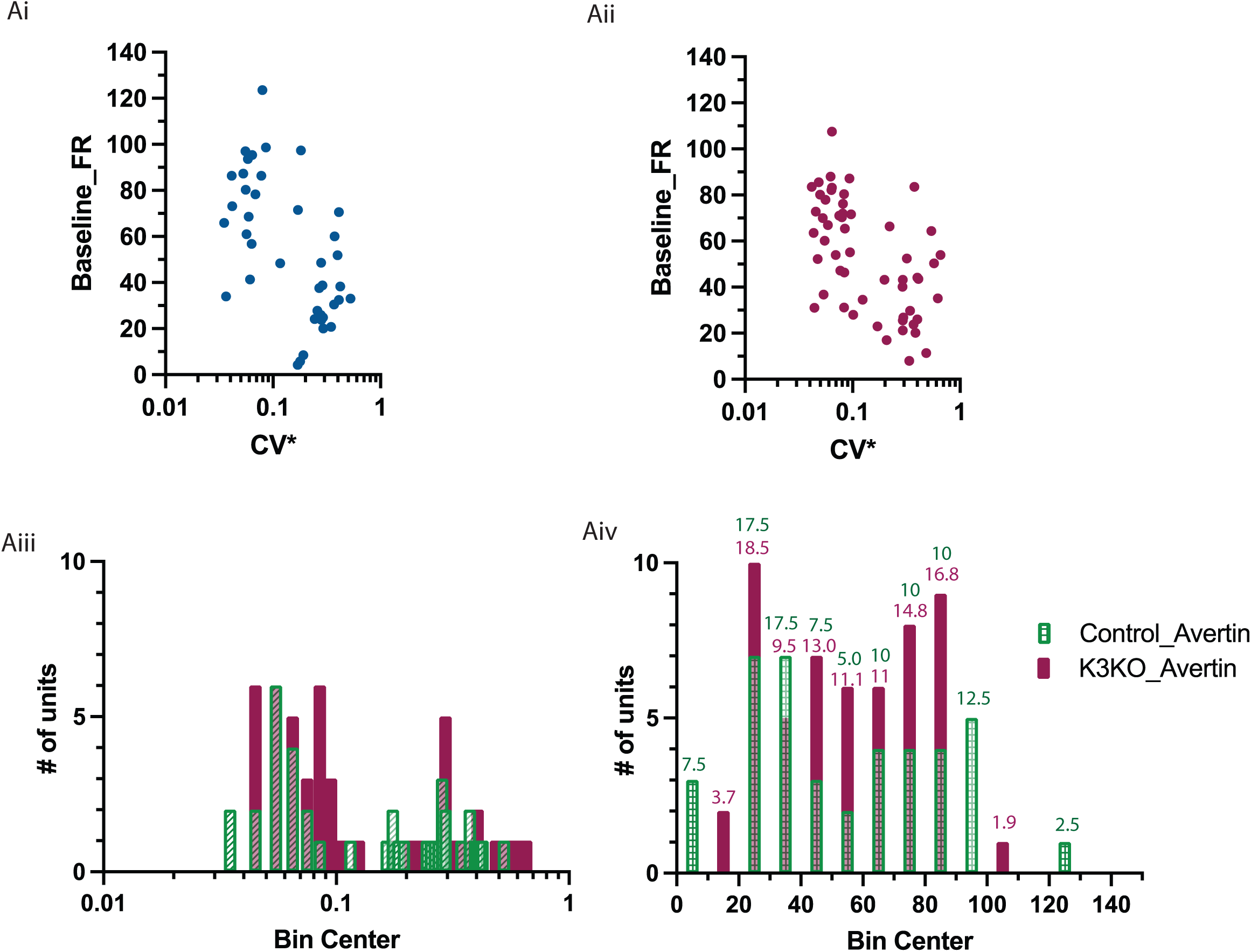
KCNQ3 knockout (KO) mice have altered spontaneous firing rate distribution of vestibular afferents. A, B, vestibular afferent discharge rate versus CV* is altered in KCNQ3 KO mice (B, red) compared to control (A, blue). C, frequency histogram showing afferent CV* in controls (green) vs KCNQ3 KO mice (red). D, frequency histogram showing afferent baseline discharge rate for control (green) vs KCNQ3 KO mice (red) with numbers on top representing %unit in that bin.

### KCNQ3 KO mice have altered EVS-mediated slow response in regular afferents but not irregular firing afferents

Next, we explored whether EVS-mediated slow excitation in vestibular afferents of KCNQ3 KO mice have been altered. Similar percentage of regular and irregular units were recorded control and KCNQ3 KO mice groups (Control vs KCNQ3 KO: Regular: 43.4% vs 46.8%; Irregular: 56.6% vs 53.2%). In both control and KCNQ3 KO, the amplitude of slow response increased as a function of CV* (Figs. 6A, 6B). However, the slope of the linear regression line was significantly different with slope Y=59.10*X + 5.785 for control and Y = 34.90*X + 12.38 for KCNQ3 KO (F(1, 115) = 4.636, p=0.0334) (slow response amplitude <5sp/s were excluded from the analysis to improve signal to noise ratio) (Fig. 6C). The slow response amplitude was significantly different between the groups in regular firing afferents but not in irregular firing afferents (Figs. 6E, 6F) (regular: 8.52±0.58 vs 14.16±1.65, t(29.44)=3.252, p=0.0031, unpaired Welch’s t-test; irregular: 23.92±2.22 vs 24.83±2.93, t(61)=0.2452, p=0.8072, Unpaired student’s t-test). Figure 6D shows the average slow response amplitude for each bin for the CV* (Fig. 6D) (bin size: for CV*<0.1, afferents were binned into three groups (CV*<0.045, <0.075, <0.1) and CV*>0.1 afferents were binned into incremental bin size of CV* of 0.5).

**Figure 6.**
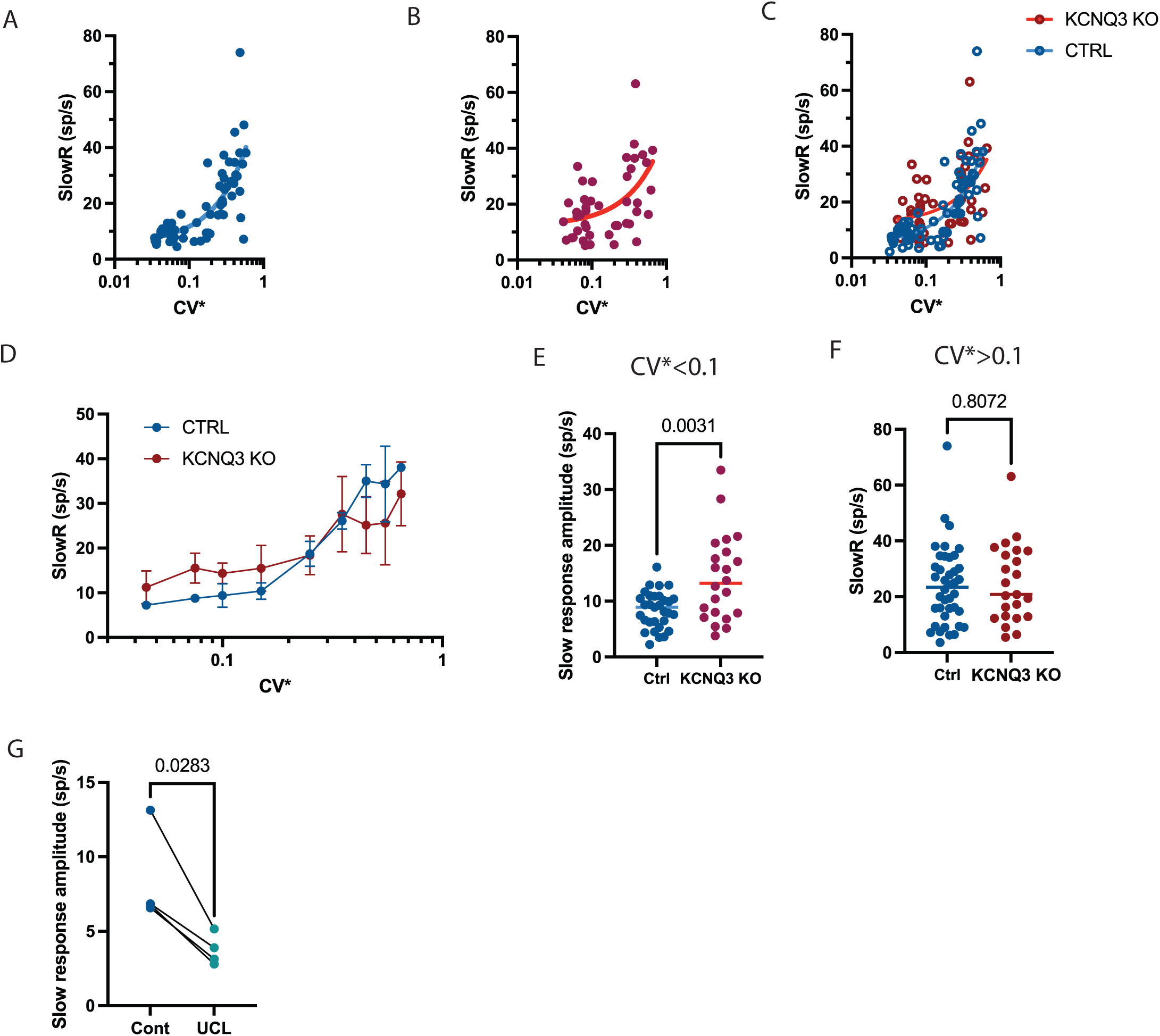
KCNQ3 knockout (KO) mice have larger slow response in regular firing afferents: A, In control WT (A, blue) (40 units from 6 mice, SR>5sp/s) mice, distribution of slow response to CV* has line of best fit of y=65.22x + 5.086 with r2 of 0.77. Whereas KCNQ3 KO (B, red) (54 units from 7mice, SR>5sp/s), distribution of slow response to CV* has line of best fit of y=34.90x + 12.38 with r2 of 0.25. C, superimposed slow response distributions. D, Average slow response amplitude across CV* (bin length: regular=0.01; irregular=0.1). E, Slow response in regular afferents (CV*<0.1) in control (CTRL, blue) vs KCNQ3 KO (K3KO, red) mice. F, Slow response in irregular afferents (CV*>0.1) in control (CTRL, blue) vs KCNQ3 KO (K3KO, red) mice. G, I. Representative continuous rate histogram of an irregular afferent (G) and a regular afferent (I) in KCNQ3 KO post-administration of 20mg/kg of UCL. H, In KCNQ3 KO mice post-administration of 20mg/kg of UCL. KCNQ2 blocker diminishes 54% of slow response in regular firing afferents followed by enhancement in firing afferents in KCNQ3 KO mice.

This was opposite to what we had hypothesized as we expected mice lacking the KCNQ3 channel subunit will have diminished EVS-mediated slow response in regular firing afferents. To reconcile these observations with our pharmacological data, we next explored the KCNQ channel subunit responsible for the presumed compensation of the EVS-mediated slow response in regular firing afferents in KCNQ3 KO mice.

### KCNQ2 contributes to slow excitation in regular afferents in KCNQ3 KO mice

Our initial hypothesis was that contribution of KCNQ2 in EVS-mediated slow excitation in regular afferents in KCNQ3 KO mice will be larger compared to control. IB administration of the KCNQ2 blocker UCL reduced 53.71% of slow response in KCNQ3 KO mice (Fig. 6G) (CV*<0.2: 8.32±1.6 vs 3.74±0.55, t(3)=3.985, p=0.0283, Paired student’s t-test). This suggests that in absence of the KCNQ3 subunit, KCNQ2 compensates for part of the EVS-mediated slow response. We also observed a fall in baseline followed by enhancement in EVS-mediated slow excitation (data not shown).

### VsEP in KCNQ3 KO mice

KCNQ3 KO mice had altered a resting discharge rate distribution versus afferent CV*. To test if the altered resting discharge rate affected the functionality of the afferents, we measured vestibular sensory-evoked potentials (VsEPs), which is a population response from irregular firing afferents to transient changes in linear head acceleration (Curthoys et al., 2017; Jones et al., 2015; Lee et al., 2017; Ono et al., 2020). The peripheral P1N1 amplitude was not significantly different between the control (n=9) and KCNQ3 KO mice (n=8-11) (Fig. 7A, F (1,26) =0.1903, p=0.6662). However, KCNQ3 KO animals had lower P2N1 amplitudes compared to control mice, although the difference was not statistically significant (Fig. 7B, F (1, 26) = 0.7358, p=0.3988). The P1 latency was significantly higher in KCNQ3 KO compared to control (Fig. 7C, P1: F (1,26) =7.5140, p=0.0109). However, the N1 latency did not show a significant difference between the two groups (Fig. 7D, F (1,26) =2.787, p=0.1070). These observations suggest that the loss of KCNQ3 is likely altering central vestibular circuitry.

**Figure 7.**
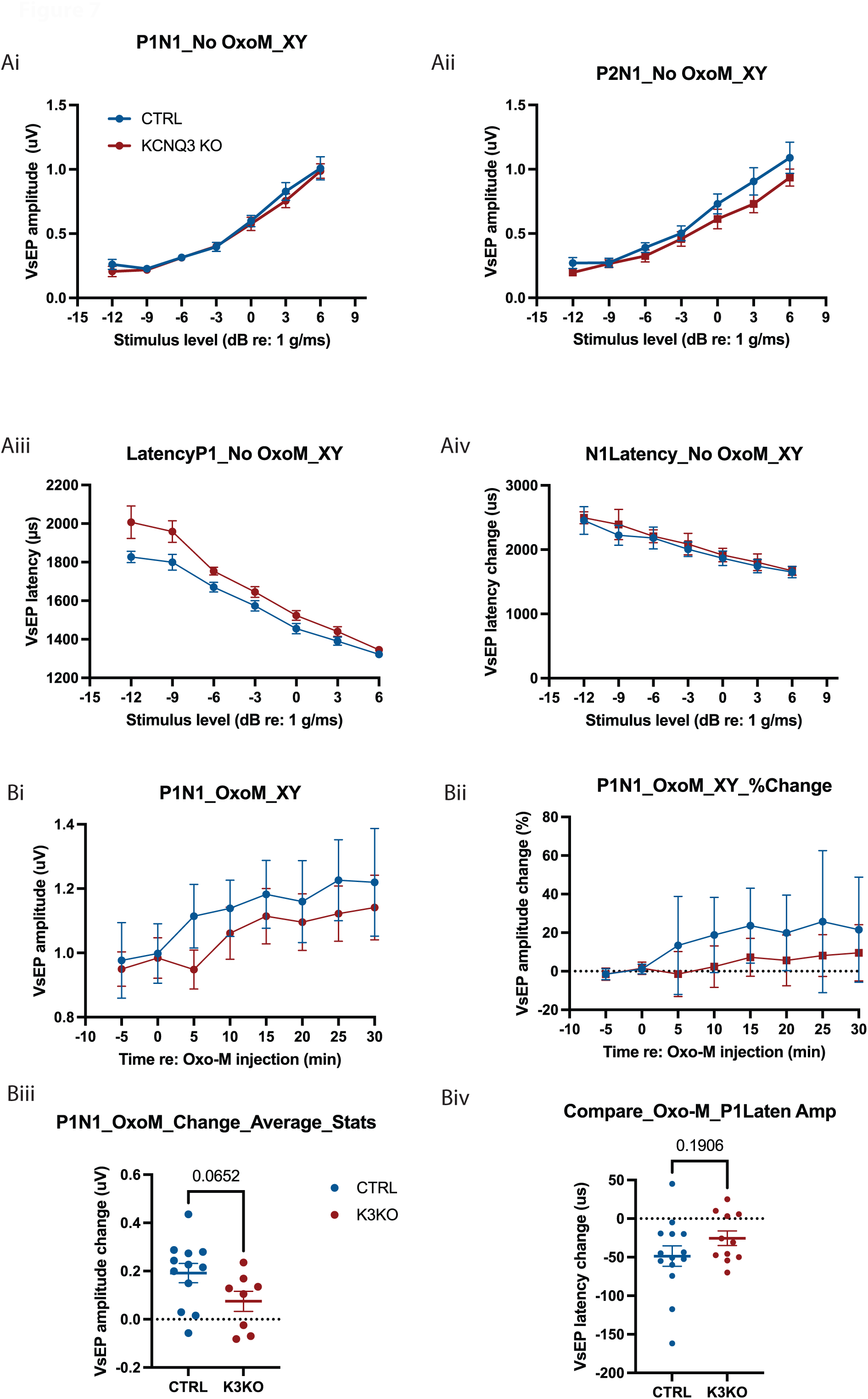
KCNQ3 knockout (KO) mice have altered VsEP. A-D, VsEP measured in control (blue, CTRL) and KCNQ3 KO (red, K3KO). A, P1N1 amplitude in CTRL (blue) vs KCNQ3 KO (red) mice at stimulus intensity from –12 dB re: 1.0 g/ms to +6 dB re: 1.0 g/ms, (F (1,26) =0.1903, p=0.6662). B, P2N1 amplitude in CTRL (blue) vs KCNQ3 KO (red) mice at stimulus intensity from –12 dB re: 1.0 g/ms to +6 dB re: 1.0 g/ms, (F (1, 26) = 0.7358, p=0.3988). C, P1 latency in CTRL (blue) vs KCNQ3 KO (red) mice at stimulus intensity from –12 dB re: 1.0 g/ms to +6 dB re: 1.0 g/ms, (F (1,26) =7.5140, p=0.0109). D, N1 latency in CTRL (blue) vs KCNQ3 KO (red) mice at stimulus intensity from –12 dB re: 1.0 g/ms to +6 dB re: 1.0 g/ms, (F (1,26) =2.787, p=0.1070). E-H, Effect of Oxo-M on VsEP amplitude and latency in KCNQ3 KO mice. E, P1N1 amplitude post-administration of Oxo-M. F, Normalized VsEP P1N1 amplitude (% Change) over a period of 30mins following Oxo-M administration in control and KCNQ3 KO. G, Average P1N1 amplitude at time 10-30mins post-Oxo-M administration in control (blue, CTRL) and KCNQ3 KO (red, K3KO). H, Average P1 latency at time=15mins post-Oxo-M administration in control (blue, CTRL) and KCNQ3 KO (red, K3KO).

We next wanted to explore the effect of mAChR activation of on VsEP metrics in KCNQ3 KO mice. Previous research (Sinha et al., 2023) demonstrated that the activation of mAChRs using Oxotremorine-M (Oxo-M), a mAChR agonist, enhanced VsEP P1N1 amplitude and reduced P1 latency in control mice (Sinha et al., 2023). Here, we repeated this experiment in KCNQ3-KO mice.

We measured VsEP waveforms at +6 dB re: 1.0 g/ms before and after the IP administration of 2mM Oxo-M. Consistent with previous findings (Sinha et al., 2023), we observed an increase in P1N1 amplitude over time in both control and KCNQ3 KO mice following Oxo-M administration (Fig. 7E). To quantify the changes, we analyzed the percentage change in P1N1 amplitude, with time, relative to the respective pre-Oxo-M condition in both groups. The percentage enhancement in P1N1 amplitude was significantly greater in control mice compared to KCNQ3 KO mice (Fig. 7F, F (1,23) =9.712, p=0.0049). However, the average change in P1N1 amplitude after 10 mins post-Oxo-M, between the groups, was not significantly different (Fig. 7G, 0.1917±0.03976 us vs 0.07479±0.04170us, t(18)=1.963, p=0.0652). The P1 latency was not significantly different between control and KCNQ3 KO (Fig. 7H) (–48.54±13.17us vs –25.45±9.512us, t(23)=1.348, p=0.1906, one-way ANOVA with Tukey’s multiple comparison).

## Discussion

Various KCNQ channel subunits are present in different microdomains of calyx-bearing afferent terminals (Hurley et al., 2006; Lysakowski et al., 2011). In this study, we explored which KCNQ channel subunits might be modulated by electrical stimulation of EVS neurons and can contribute to EVS-mediated slow excitation in mammals. Previous work in turtle and rat suggested that KCNQ channels are the downstream targets of mAChR activation given their sensitivity to the KNCQ blockers, linopirdine and XE991 and lack of effect from the ERG channel blockers, E-4031 (Holt et al., 2017; Perez et al., 2009; Perez et al., 2010; Ramakrishna et al., 2021; Sinha et al., 2023). In this paper, we confirmed a similar role for KCNQ channels in our *in vivo* mouse preparation. We show that EVS-mediated slow excitation of vestibular afferents is blocked by XE991 but not E-4031. And, in most cases, application of XE991 enhanced the baseline firing rate of afferents, consistent with closing of channels.

In this study, the KCNQ2/3 opener ICA113081 resulted in an enhancement of the EVS-mediated slow response, similar to the observations seen with retigabine. KCNQ3 is present on the outer surface of dimorphic endings and down along their afferent trunk towards the heminode. The large enhancement in EVS-mediated excitation of regular afferents suggests that KCNQ3 channels in the two domains might be both targeted by EVS neurons. Blockade of EVS-mediated excitation by the KCNQ2 channel blocker, UCL2077, which can also have some inhibitory effect on KCNQ2/3 heteromeric channels, was larger in irregular afferents compared to regular afferents. This suggests that the KCNQ2 channel subunit is likely a key mediator of slow response in irregular afferents. It is worth noting that KCNQ2/3, KCNQ2/5, KCNQ3/4, KCNQ3/5, and KCNQ4/5 heteromeric channels have also been found in mouse (Bal et al., 2008; Lerche et al., 2000; Schroeder et al., 2000; Schroeder et al., 1998; Søgaard et al., 2006; Soh et al., 2022; Wang et al., 1998). Additionally, KCNQ2/3/5 trimeric channels can also be formed in vitro (Soh et al., 2022). While the presence of multiple KCNQ (KCNQ2-5) channel subunits complicate some of the interpretation of our observations, such heterogeneity, however, might explain the differential blockade of KCNQ channels among different vestibular afferent populations. KCNQ2/3 openers lead to enhancement in slow excitation in most of the regular afferents while KCNQ2 blocker blocked slow excitation in irregular afferents. Additionally, KCNQ2/4 opener led to enhancement in few of the afferents. Moreover, both KCNQ2/3 and KCNQ2/4 opener led to decrease in baseline firing rate of vestibular afferents, while KCNQ2 blocker did not affect the baseline firing rate. Our results suggest that KCNQ3 and KCNQ4 contributes to resting discharge rate of vestibular afferents. Moreover, KCNQ2 and KCNQ3 subunits are likely key mediators of M-current in irregular afferents and regular afferents, respectively.

Unlike KCNQ2 and KCNQ3 subunits, which are expressed on the inner surface or on the outer surface of the calyx endings, respectively, the KCNQ4 subunit is expressed on various microdomains on both the inner and outer face which differs between dimorphic and calyx-only afferents (Hurley et al., 2006; Lysakowski et al., 2011). This is consistent with the variation we saw in the effect of the KCNQ2/4 opener, ML213, on EVS-mediated slow excitation. In most of the irregular afferents, administration of ML213 led to a gradual decline in baseline firing rate. This is consistent with presence of KCNQ4 channel subunits on the heminode in central calyces and at a lower concentration on the outer surface of calyx endings. In the regular afferents, however, the decline in afferent firing rate was slower followed by an abrupt fall in afferent firing rate. This would suggest that in the regular afferents, the KCNQ4 channel on the outer surface is at a lower concentration, which is affected KCNQ2/4 opener. This is in congruence with the IHC staining against KCNQ4 (Lysakowski et al., 2011). The KCNQ4 channels present on the inner surface of the calyx has been suggested to important for the maintenance afferent excitability as well as to facilitating cross-talk between type I HCs and calyx afferents. Additionally, KCNQ4 channels have been suggested to contribute to non-quantal mode of transmission (Contini et al., 2020; Contini et al., 2017; Goldberg, 1996a, 1996b; Govindaraju et al., 2023; Highstein et al., 2014; Lysakowski & Goldberg, 1997). Hence, affecting KCNQ4 subunits on the inner surface can disrupt the resting discharge rate of afferents as well. In few units, the KCNQ2/4 opener showed enhancement in the amplitude of EVS mediated slow response. This could be an effect of action of the opener on KCNQ2 channel subunit or due to KCNQ4 present on the heminode in calyx-only afferents and on the outer surface of afferents (Lysakowski et al., 2011). KCNQ4 subunits on the outer surface and heminode can be directly modulated by EVS neurons as a function of their close proximity to efferent endings. However, KCNQ4 subunits might be contributing to EVS-mediated slow response but due to technical limitation of our preparation we were not able to distinguish the afferent type (beyond regularity) where the KCNQ2 subunit contributed to the EVS-mediated slow response.

Alternatively, the effects of UCL2077 on EVS-mediated slow response in irregular afferents might originate from block of KCNQ4 on the inner surface of calyx endings. However, this seems unlikely as there is no change in baseline firing of afferents. Due to the high concentration of KCNQ4 channels on the inner surface, compared to other KCNQ subunits, blocking of KCNQ4 channels should increase the baseline firing rate which we did not observe in our preparation. This suggests that the effect we saw with application of UCL2077 is due to blocking of KCNQ2 channels on the inner surface.

We were not able to evaluate role of KCNQ5 subunit due to lack of selective drugs against KCNQ5 subunit. However, UCL2077 acts as an agonist for KCNQ5 channels. But we did not see any enhancement in presence of UCL2077 in wildtype mice. Although it is possible that the enhancement was obscured by the inhibition of slow response when blocking the KCNQ2 channels. Intriguingly, no change in the baseline firing rate of the afferent was observed in the control even after closing of the KCNQ2 channels. This might arise from the fact that UCL enhanced the KCNQ5 channel activity while blocking the KCNQ2 activity (Soh & Tzingounis, 2010). Both KCNQ2 and KCNQ5 are present on the inner surface of calyx afferent endings (Lysakowski et al., 2011).

However, in KCNQ3 KO mice, UCL 2077 showed inhibition of EVS-mediated slow response accompanied with fall in afferent firing rate. The fall in baseline firing rate was followed by enhancement in slow response. This suggests that KCNQ5, for which UCL2077 acts as an agonist, might have compensated for KCNQ3 subunit in KCNQ3 KO mice, in addition to KCNQ2. Recently, (Soh et al., 2022) showed that in KCNQ3 KO mice, KCNQ2 subunit forms heteromeric channels with KCNQ5 subunit, in cortex and hippocampus, but not in WT mice (Soh et al., 2022). Hence, KCNQ2 subunit with KCNQ5 subunit might mediate M-currents in vestibular afferents in KCNQ3 KO mice. However, a clear role of KCNQ5 in WT animal is yet to be explored.

In vestibular afferents, KCNQ subunits are present on both the inner and outer surface of the calyx ending (Lysakowski et al., 2011). Efferent endings synapse on the outer surface of the calyx endings and can activate the mAChRs present in close proximity to the terminal (Holt et al., 2017). The activated mAChR, however, can close the KCNQ channels present both on the outer surface and on the inner surface. Activation of mAChRs can presumably close KCNQ channels in various microdomain through varied secondary messengers. Activated mAChR activates membrane delimited phospholipase Cβ (PLCβ), which hydrolyses phosphatidylinositol-4,5-bisphosphate (PIP2) in the membrane. PIP2 is required for KCNQ channels to stay open (Suh & Hille, 2002) and hydrolysis of PIP2 leads to closure of KCNQ channels. This mechanism will close KCNQ channels in close proximity to the activated mAChR. Other secondary messengers can target channels that are further away from the site of mAChR activation. PLCβ hydrolyses PIP2 into inositol-1,4,5-triphosphate (IP3) and diacylglycerol (DAG). IP3 can diffuse in the cell and can bind to IP3R on the ER. This opens the IP3R on the ER and increase the concentration of intracellular Ca2+. Calmodulin (CaM) is bound to open KCNQ channels. CaM is sensitive to intracellular Ca2+. Binding of Ca2+ to CaM decreases the affinity of CaM to KCNQ channel and KCNQ undergoes confirmational changes which closes the channels ((Gamper & Shapiro, 2003), (Gamper & Shapiro, 2007), (Chambard & Ashmore, 2005), (Kosenko & Hoshi, 2013), (Sihn et al., 2016)). This mechanism can lead to closure of the KCNQ channels that are further away from activation site.

Interestingly, in control mice, slow response amplitude increases as a function of CV*. However, in KCNQ3 KO mice, this relation was disrupted. This suggests that the difference in the amplitude in EVS-mediated slow response between regular and irregular afferents, might be a function of the KCNQ channel subunit involved, and the location of these subunits on the afferents. Additionally, in KCNQ3 KO mice, KCNQ2 and possibly KCNQ5 might be main regulator of EVS-mediated slow response in afferents.

In conclusion, this study attempted to explore contribution of various KCNQ channel subunits to resting discharge rate of primary vestibular afferents and to the EVS-mediated slow response. Here, we show that KCNQ2/3 was key mediator of EVS-mediated slow response and KCNQ3/4 are the key channels involved in maintaining resting discharge rate in primary vestibular afferents.

## Acknowledgment

This research was supported by NIH/NIDCD Grants R01DC0016974 (JH). We would like to thank Dr. Thomas J. Jentsch at Leibniz-Forschungsinstitut für Molekulare Pharmakologie (FMP) and Max-Delbrück-Centrum für Molekulare Medizin (MDC), Berlin, Germany, for providing us with KCNQ3 knockout mice.

